# Automated ensemble assembly and validation of microbial genomes

**DOI:** 10.1101/002469

**Authors:** Sergey Koren, Todd J. Treangen, Christopher M. Hill, Mihai Pop, Adam M. Phillippy

**Author notes:** Corresponding author Email addresses: SK TJT CMH MP AMP.

## Abstract

**Background:** The continued democratization of DNA sequencing has sparked a new wave of development of genome assembly and assembly validation methods. As individual research labs, rather than centralized centers, begin to sequence the majority of new genomes, it is important to establish best practices for genome assembly. However, recent evaluations such as GAGE and the Assemblathon have concluded that there is no single best approach to genome assembly. Instead, it is preferable to generate multiple assemblies and validate them to determine which is most useful for the desired analysis; this is a labor-intensive process that is often impossible or unfeasible.

**Results:** To encourage best practices supported by the community, we present iMetAMOS, an automated *ensemble assembly pipeline*; iMetAMOS encapsulates the process of running, validating, and selecting a single assembly from multiple assemblies. iMetAMOS packages several leading open-source tools into a single binary that automates parameter selection and execution of multiple assemblers, scores the resulting assemblies based on multiple validation metrics, and annotates the assemblies for genes and contaminants. We demonstrate the utility of the ensemble process on 225 previously unassembled *Mycobacterium tuberculosis* genomes as well as a *Rhodobacter sphaeroides* benchmark dataset. On these real data, iMetAMOS reliably produces validated assemblies and identifies potential contamination without user intervention. In addition, intelligent parameter selection produces assemblies of *R. sphaeroides* that exceed the quality of those from the GAGE-B evaluation, affecting the relative ranking of some assemblers.

**Conclusions:** Ensemble assembly with iMetAMOS provides users with multiple, validated assemblies for each genome. Although computationally limited to small or mid-sized genomes, this approach is the most effective and reproducible means for generating high-quality assemblies and enables users to select an assembly best tailored to their specific needs.

## Background

Genome assembly reconstructs a genome from many shorter sequencing reads as faithfully as possible [1–3]. Since reasonable formulations of the problem are NP-hard [2, 4], practical implementations often return an approximate solution that contains errors. Recent assembly evaluations like GAGE and the Assemblathon [5–8] have highlighted the chaotic nature of genome assembly, in which assembler performance varies widely across datasets and small parameter changes can have drastic effects. In GAGE-B [8] for example, each dataset required a different k-mer parameter, the best assembler was not consistent across datasets, and the continuity difference between best and second best was often two-fold.

Although genome assembly is a complex problem, validating assemblies is more straightforward. The quality of a genome assembly can be confirmed by verifying that the layout of reads is consistent with the sequencing process used to generate the data [9]. Multiple tools have been recently developed for validating genome assemblies both with and without the use of a reference genome [10–15].

Thus, given the chaotic nature of assemblers and the relative ease of validation, it is recommended to generate multiple assemblies and use validation to determine the most appropriate one. This is akin to a “hypothesis generation” view of assembly [16], which can be most easily implemented as an ensemble of independent methods. Unfortunately, running multiple assemblers is a time consuming, non-trivial task requiring substantial installation, learning, and maintenance costs.

There exist a limited set of tools that integrate automated parameter selection and validation into the assembly process. The A5 pipeline [17, 18] automates the microbial assembly process, but is limited to a single assembler and includes limited validation. CG-Pipeline [19] is targeted to 454 sequencing. VelvetOptimizer [20] automates a parameter sweep of k-mer sizes for the Velvet assembler [21], but uses contig N50 size as the optimization metric, which is not always representative of assembly quality [7, 22]. More recently, a number of assembly methods have been developed that incorporate assembly likelihood estimates into the primary assembly algorithm [23–25]. However, none of these tools robustly automate the execution of multiple assembly methods and validation metrics to achieve the best possible assembly. Here we present iMetAMOS, which automates the process of ensemble assembly and validation.

## Implementation

Whereas MetAMOS [26] was developed for metagenomic assembly, iMetAMOS is an isolate-focused extension that encapsulates the current best practices for microbial genome assembly using Illumina [27], 454 [28], Ion [29], or PacBio [30] sequencing data. Building on the conclusions of GAGE and the Assemblathon, iMetAMOS runs multiple, independent tools to generate and validate assemblies. Uniquely, iMetAMOS automates the entire ensemble assembly process including automated parameter selection and sweeps, execution of multiple assembly and validation tools, preliminary gene annotation, and identification of potential contaminating sequences. This ensemble approach is robust to individual tool failures and reliably generates high-quality assemblies with minimal user input.

### Pipeline Design

Figure 1 details the iMetAMOS pipeline, which treats each assembly project as a competition among multiple assemblers. As an added benefit to the ensemble approach, iMetAMOS is robust to failure; if any tool or parameter combination fails, iMetAMOS will continue the analysis using only the ones that succeeded. A “winning” assembly is then automatically selected based on the combined validation results. However, users may also browse the validation metrics to choose their own preferred assembly, such as one that optimizes consensus accuracy without regard to continuity.

**Figure 1.**
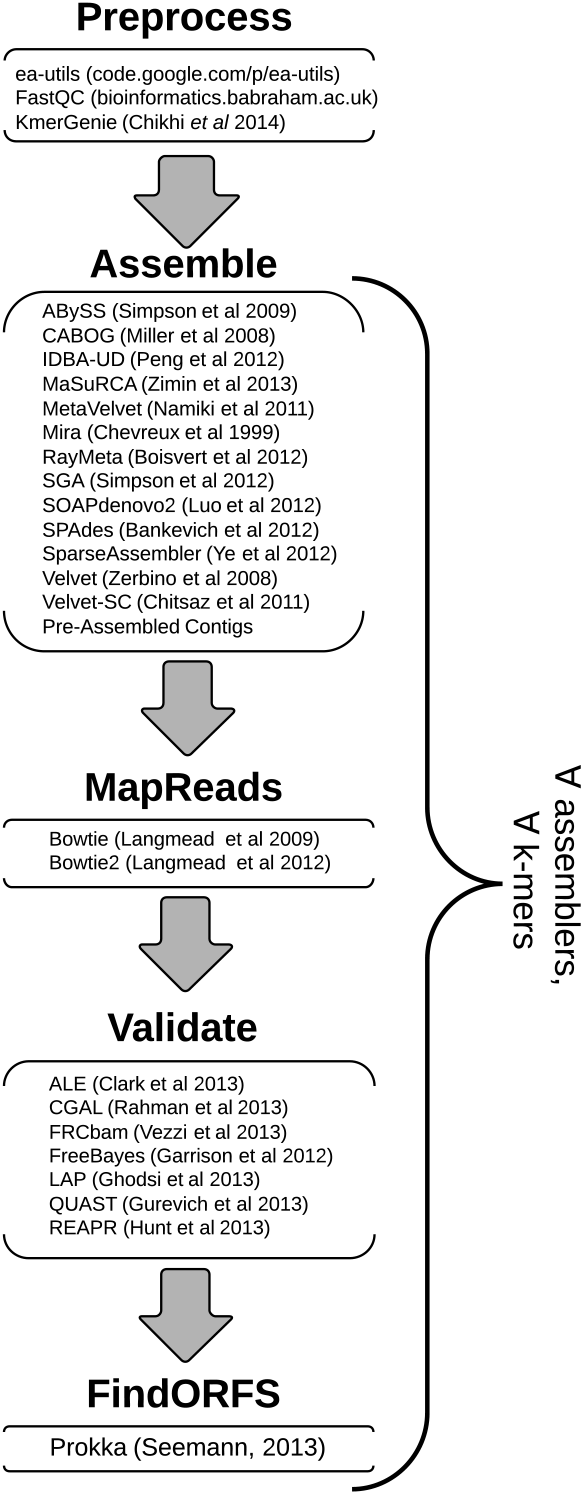
iMetAMOS workflow and incorporated tools. iMetAMOS currently incorporates 13 assemblers [21, 33–38, 40–45] and 7 validation tools [10–15, 47]. Prokka [50] is used to predict genes and annotate all assembiles. Users can control the suite of assemblers and validation tools to be executed, as well as the scoring formula used to choose the best assembly. This assembly is evaluated for the presence of contamination.

iMetAMOS is primarily written in Python and builds upon Ruffus [31] for pipeline management. However, it incorporates many freely available tools written in a variety of languages. To simplify installation, iMetAMOS is distributed as 64-bit OS X and Linux binaries, including all supported assemblers, tools, and required databases. On 32-bit systems, iMetAMOS automatically downloads and installs the required dependencies, as needed, which significantly simplifies installation.

### Modular Infrastructure

To support future extensibility, iMetAMOS includes a generic framework to add new tools to the pipeline. Currently supported modules are for assembly and classification.

When a new tool is available, no code modification is need. Instead, a configuration is written to specify parameters for the tool and required inputs and outputs. iMetAMOS will automatically load this configuration and run the requested tool. When an external tool is executed, a corresponding citation is output to ensure users of iMetAMOS properly credit the tools on which it relies.

### Reproducibility

iMetAMOS enables reproducible analysis by recording all commands, software versions (via an MD5 hash), and intermediate inputs and outputs. The single, comprehensive binary is generated via PyInstaller [32], which also serves to fix and archive the exact version of all programs used. Reproducibility of custom analyses is supported via workflows. A workflow defines the software required for an analysis, as well as optional parameters and input data. Workflows support both local and remote file names, as well as SRA run identifiers, and can inherit their parameters from other workflows, allowing users to easily add or modify input data or parameters. Given a workflow, iMetAMOS will download any required remote data and run the analysis using pre-specified parameters. For further reproducibility, a workflow is automatically created for every iMetAMOS run, which can be easily shared with remote collaborators. If the data are available on the Internet, the entire analysis can be reproduced by two simple commands.

### Assembly

Assembly is treated as a hypothesis generation and testing problem. Multiple assembly tools are run to ensure robustness to failure and a thorough exploration of the hypothesis space. The following assemblers are currently supported: ABySS [33], CABOG [34], IDBA-UD [35], MaSuRCA [36], MetaVelvet [37], MIRA [38], Ray/RayMeta [39, 40], SGA [41], SOAPdenovo2 [42], SPAdes [43], SparseAssembler [44], Velvet [21], and Velvet-SC [45]. For De Bruijn assemblers, a k-mer size is automatically selected using KmerGenie [46]. Alternatively, users can specify a list of k-mers and iMetAMOS will run each assembly with each specified k-mer. In this mode, iMetAMOS can operate similarly to VelvetOptimizer [20], but for multiple assemblers and with more appropriate validation measures.

### Validation and Annotation

Each assembly is treated as a hypothesis subject to validation. The following validation tools are supported: ALE [10], CGAL [11], FRCbam [15], FreeBayes [47], LAP [14], QUAST [13], and REAPR [12]. Both reference-based and reference-free validations are performed. For reference-based validation, a MUMi distance [48] is used to recruit the most similar reference genome from RefSeq [49] to calculate reference-based metrics. For reference-free validation, the input reads and read pairs are verified to be in agreement with the resulting assembly using both likelihood-based methods and mis-assembly features. In addition, to provide an initial annotation and comparison between gene content, the assemblies are automatically annotated using Prokka [50].

From the ensemble, the “winning” assembly is selected using the consensus of the validation tools. For each selected metric, the assemblies are assigned an order from best to worst (with 1 being best). By default, the top assemblies are selected as those that are in the top 10% for at least half the metrics. The best assembly is then selected as the top scoring assembly with the highest count of best scores. A user can select a single metric (i.e. consensus accuracy) or an arbitrarily weighted combination of metrics for validation. Importantly, this allows users to customize the validation process to suit their downstream project goals. For example, studies focused on phylogenetic tree reconstruction may prefer to minimize consensus errors, while structural variation studies may instead focus on maximizing continuity and minimizing long-range errors.

### Contamination Detection

Although iMetAMOS focuses on single-genome assembly, all inputs are considered as a metagenome to control against possible contamination. The winning assembly’s contigs and unassembled reads are analyzed by a taxonomic classification program. By default, iMetAMOS uses the k-mer based Kraken [51] tool, but the alternative methods of FCP [52], PhyloSift [53], PHMMER [54], and PhymmBL [55] are also supported. Contigs are partitioned into separate, taxon-specific directories (genus by default) according to their classification, so that contaminating sequence can be easily identified and removed. This process also serves as an initial species identification when assembling novel organisms.

The classification result is dependent on the classifier and database used, and serves as only a preliminary species identification or indicator of potential contamination. Manual follow-up is recommended to confirm the classification. For example, recently acquired genomic elements, such as phage integrations, may be incorrectly classified. Nevertheless, this initial binning facilitates rapid identification of the assembled organism and easier contaminant removal before downstream analysis or submission to a nucleotide archive.

### Results Display

The final output of iMetAMOS is a self-contained HTML5 summary page. Here, users can browse the output files as well as drill down to detailed results from any step in the pipeline. This includes FastQC [56] reports for the preprocess step, QUAST [13] graphs and metrics from the validation step, and an interactive Krona [57] display of the taxonomic classifications.

## Results

### Automated Assembly Evaluation

With iMetAMOS it is possible to automatically recreate an assembler evaluation for every sample. We used iMetAMOS to perform ensemble assembly of the *Rhodobacter sphaeroides* 2.4.1 MiSeq dataset from the recent GAGE-B evaluation [8]. In addition, our automated evaluation included four additional assemblers (IDBA-UD [35], SparseAssembler [44], Velvet-SC [45], and Ray [39]) and validation metrics not utilized by GAGE-B (e.g. consensus accuracy).

The GAGE-B evaluation included assemblers run with multiple, manually selected k-mers for each assembler that ranged from a minimum of 31 to a maximum of 85. For a thorough comparison, the *R. sphaeroides* dataset was downloaded from the GAGE-B website (http://ccb.jhu.edu/gage_b/), and two iMetAMOS runs generated, both with an auto-selected *k* of 35 as well as all k-mer sizes divisible by 5 from 25–125. The original GAGE scripts [7] were used to calculate both corrected and raw N50s on all iMetAMOS assemblies and those from GAGE-B. In some cases, iMetAMOS ran a more recent assembler than GAGE-B. Repeating the experiment using the exact software versions as GAGE-B produced the same results, with the exception of SPAdes, which showed assembly improvement in version 2.5. The entire auto-selected k-mer iMetAMOS run can be reproduced using the following commands:

~~~
initPipeline –d test_gageb –W isolate_gageb
runPipeline –d test_gageb –p 16
~~~

The best assembly of this dataset, as selected by iMetAMOS, was MaSuRCA (*k* = 35), matching the GAGE-B result. However, the corrected N50 of the iMetAMOS MaSuRCA assembly increased to 139Kbp from the 120Kbp reported in GAGE-B. Similar improvements were observed for four assemblers when compared to the manually selected k-mer in GAGE-B. This improvement is the result selecting a value of *k* to maximize assembly correctness, rather than the GAGE-B approach of maximizing the uncorrected contig N50 size. In cases where iMetAMOS did not outperform the GAGE-B results, GAGE-B had utilized EA-UTILS [58] to preprocess the data. While EA-UTILS is supported by iMetAMOS, using raw sequencing data generated the best assemblies in GAGE-B, so pre-processing was disabled.

Figure 2 shows the raw and corrected N50s for all assemblers with *k* ranging from 25–125. This example illustrates the power of iMetAMOS, with each run being equivalent to a GAGE-style assembly evaluation. In addition, for 9 of 12 assemblers, the KmerGenie auto-selected *k* of 35 provided the best ratio of corrected to raw N50 contig size. Because the corrected assemblies are broken at each mis-assembly, the ratio of corrected to raw N50 size is a good indicator of error rate. Thus, these results suggest that simply running each assembler with an auto-selected value of *k* will produce high-quality assemblies without the computational expense of producing separate assemblies for multiple values of *k*.

**Figure 2.**
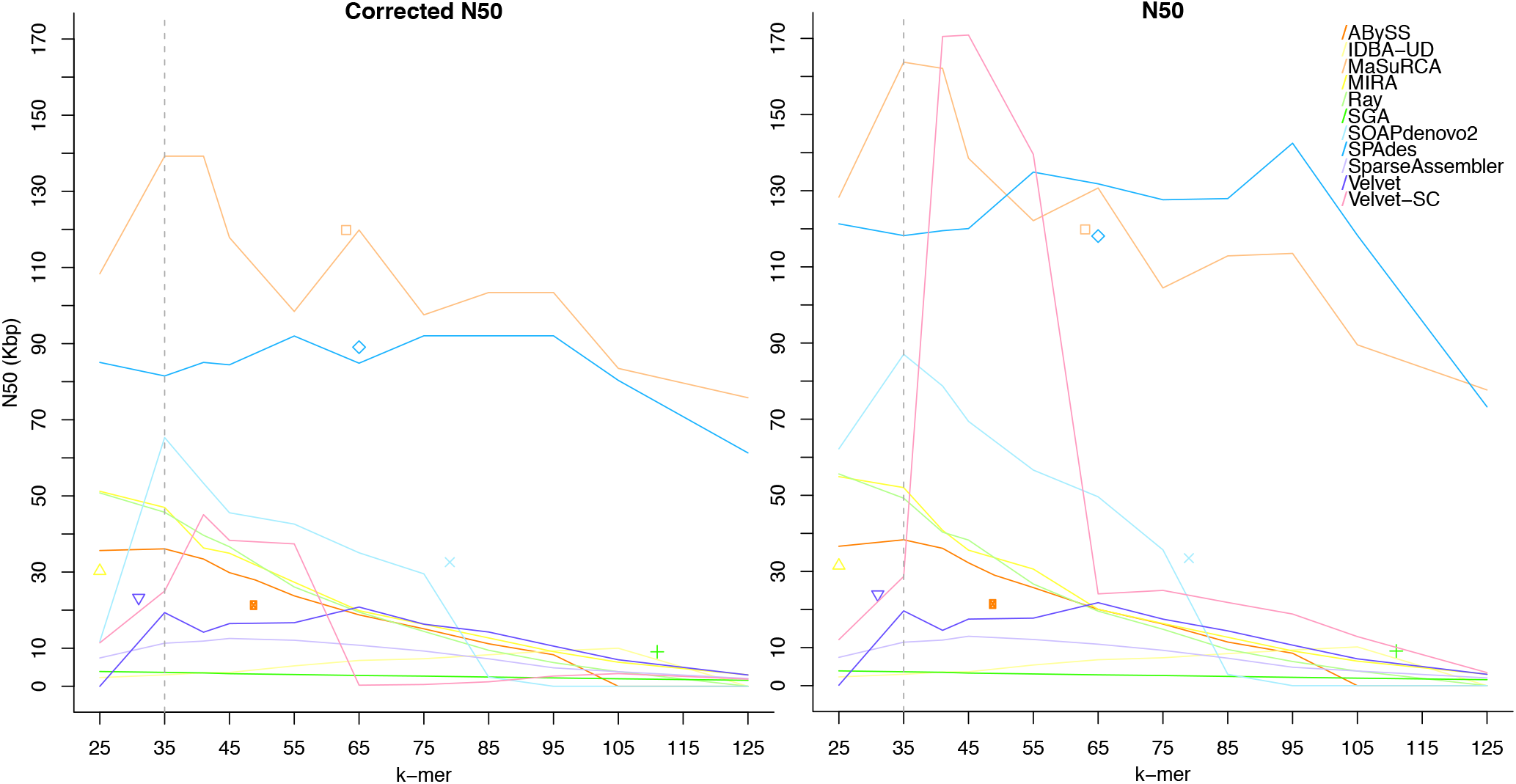
Comparison of corrected and raw N50 contig sizes for all assemblies of *R. sphaeroides*. Corrected N50 sizes were computed using the GAGE metrics [7]. The dashed vertical line indicates the auto-selected *k* of 35 chosen by KmerGenie. The individual points indicate assemblies from GAGE-B. For 9 of the 12 assemblers, the automated k-mer selection provides the best corrected to raw N50 ratio. The auto-selected k-mer also provides the best overall corrected N50 on this dataset. One notable exception is SPAdes, for which the auto-selected *k* produced an N50 13% lower than the best. This is likely caused by SPAdes use of multiple k-mers for assembly, something that KmerGenie does not currently take into account. In all other cases, except when GAGE-B used EA-UTILS [58] to trim the input sequences, the automatically selected k-mer outperforms the k-mer choice from GAGE-B.

The benefit of intelligently choosing a value of *k* in De Bruijn assembly is also obvious from Figure 2, as it changes the relative rankings of assemblies compared to the GAGE-B evaluation. In one example, MIRA goes from a corrected N50 of 15.19Kbp (ranked 6th) to 46.97Kbp (ranked 4th)—an increase of over 3-fold. The comparative ranking of assemblers on this dataset is given in Table 1.

**Table 1:**
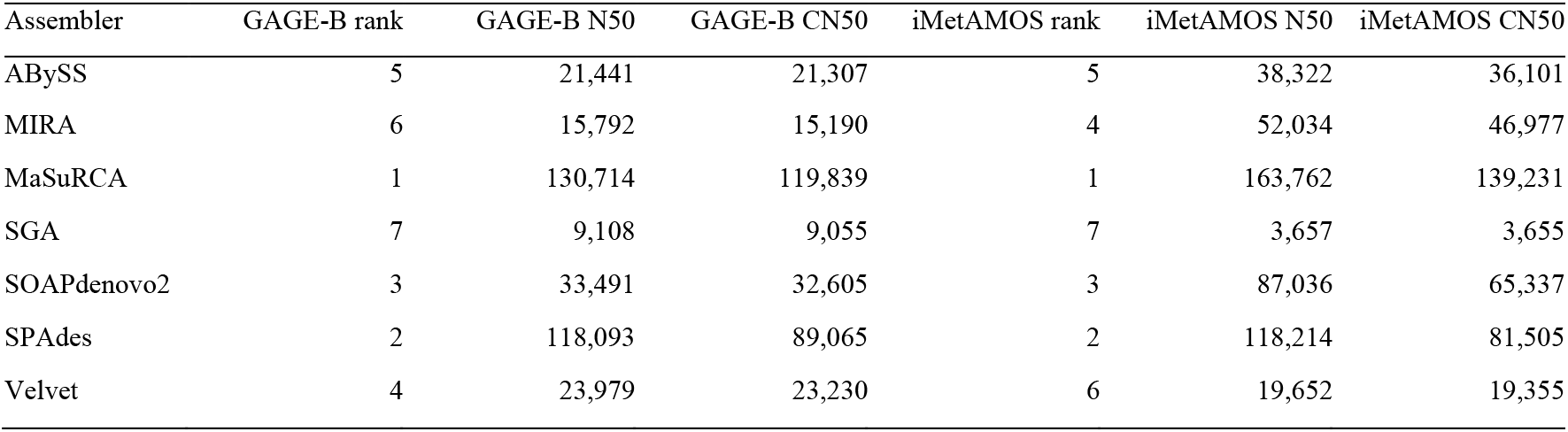
GAGE-B assemblies versus iMetAMOS assemblies on the *R. sphaeroides* dataset. The table lists the relative ranking, assembly, and corrected N50 for each of the 7 assemblers common between GAGE-B and the iMetAMOS run using an automatically-selected k-mer. Assembly N50 was computed using the reference as the true genome size. Corrected N50 (CN50) was calculated as in GAGE [7].

### Contaminant Detection

The detection and removal of contaminating DNA sequences is an often-overlooked phase of assembly. For example, the Assemblathon 1 dataset included mock contaminant, which only a few teams attempted to detect and remove [6]. Failure to remove real contaminant from assemblies significantly affects the quality of public databases to which these genomes are submitted.

To assist in contaminant detection and removal, iMetAMOS supports multiple tools that taxonomically classify the assembled contigs and reads left unassembled. To test this process on a large scale, we downloaded and analyzed the raw sequencing data from 225 samples from a recently published study of *Mycobacterium tuberculosis* [59]. All 225 runs corresponding to project ERP001731 were downloaded, and iMetAMOS was run using the ensemble of SPAdes, MaSuRCA, and Velvet. The iMetAMOS run for an individual sample can be reproduced using the following commands, which downloads the data directly from the SRA:

~~~
initPipeline –d test_TB –W isolate –m < RUN IDENTIFIER> –i
200:800
runPipeline –d test_TB –a spades,masurca,velvet –p 16
~~~

This also represents the first assembly of these samples, as the original publication focused only on read mapping (which is less affected by contamination issues). The dataset included paired-end and single-end data and sequence length ranged from 51 bp to 108 bp. The average number of contigs using unpaired data was 660, N50 was 17Kbp, and average k-mer size used was 26. For paired-end data, the average number of contigs was 333, N50 was 43Kbp, and average k-mer size used was 35. The SPAdes assembly was the best 88% of the time (Additional File 1). The full iMetAMOS analysis took an average of 32.25 hours using 16 cores. The assembly, validation, and annotation steps took approximately 50% of the runtime. Classification to identify contaminant took approximately 34% of the runtime (Additional File 2).

While the majority of samples (218 of 225) indicated no contamination, several showed signs of having multiple organisms present, such as sample ERR233356. In this case, the assembly has significantly more bases than expected (6.24 Mbp versus 4.39 Mbp) and an unusually large number of contigs (>3,700). Figure 3 shows the HTML summary for sample ERR233356, with the validation tab selected. This tab provides information on the selected assembly, quality statistics for all assemblies, as well as an indication of contamination in the sample. Figure 4 summarizes the iMetAMOS classification output for sample ERR233356. *M. tuberculosis* is clearly the dominant constituent, but a significant fraction of sequences are assigned to other bacteria or are left unclassified (because the Kraken mini classification database used includes only microbial genomes, eukaryotic contamination appears as “unclassified”). To validate the iMetAMOS output, we aligned the classified sequences to *M. tuberculosis* NITR206 (4.39 Mbp) and *S. aureus* USA300 (2.87 Mbp) using dnadiff [9]. Sequences classified as Mycobacterium had an average identity of 99.87% and 98.93% coverage of the reference. Sequences classified as Staphylococus had an average identity of 99.85% and 61.65% coverage. Over half the *S. aureus* reference genome is present in the dataset, confirming the contamination as the most likely source of these reads. We also validated the classification accuracy by comparing the reference coverage from the assembly to the reference coverage from the classified sequences. In both cases, the difference in the percentage of the reference covered by the assembly and the classified contigs was less than 1%. All 7 *M. tuberculosis* samples identified by iMetAMOS as potentially contaminanted were manually examined and had similarly large assemblies with an unusual number of sequences classified as either microbial or unknown. Using BLAST [60] confirmed both eukaryotic (human) and microbial contaminant in these samples.

**Figure 3.**
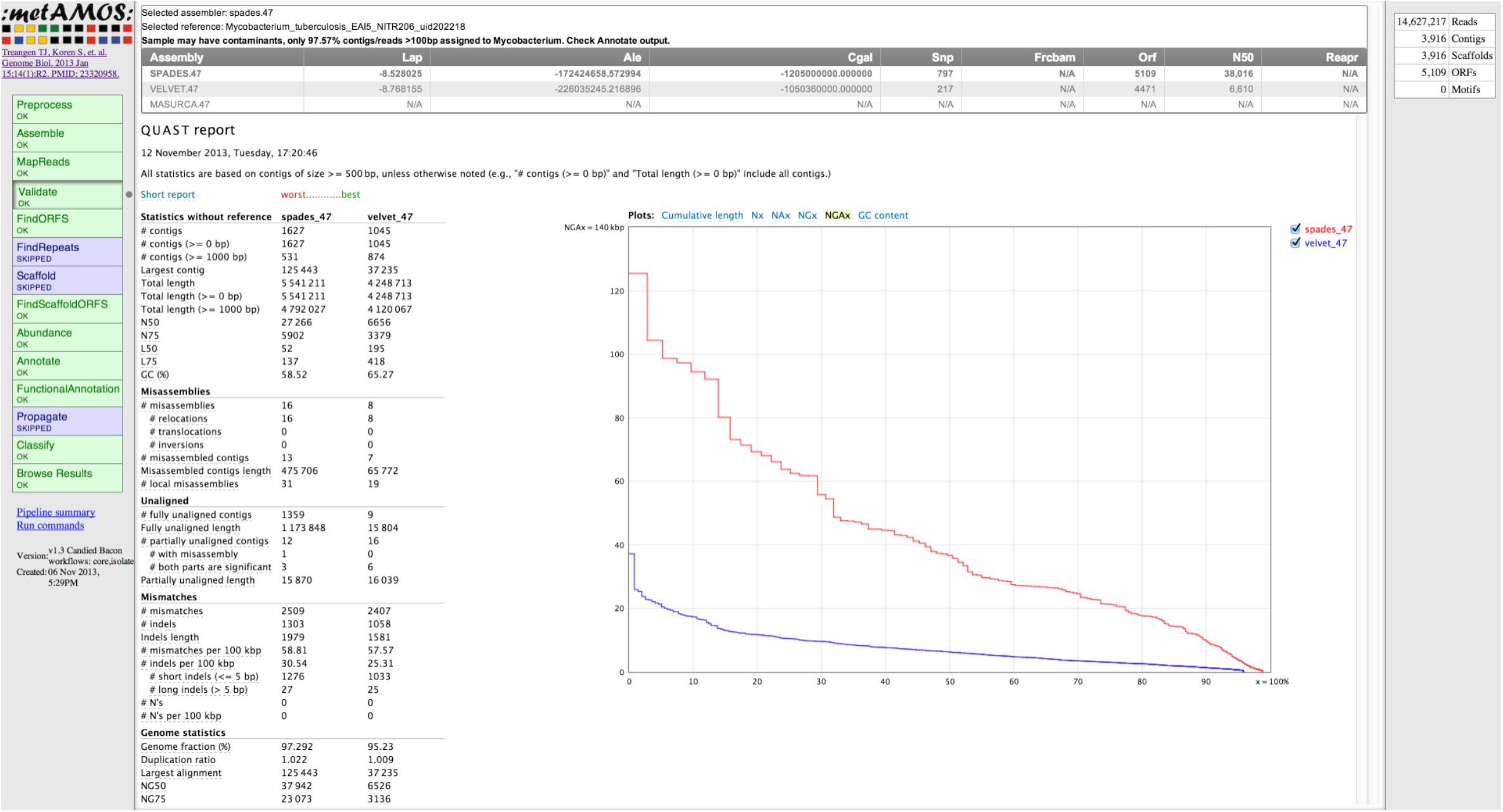
iMetAMOS validation output for an example dataset. The leftmost tab allows navigation to view the output of each pipeline step. The selected “Validation” tab results are shown in the main window. These include the validation metrics of all successful assemblies, including a comparison and QUAST [13] report against an automatically recruited reference genome from NCBI RefSeq. The validation tab also indicates the sample shows signs of contamination. Finally, the rightmost tab shows a quick summary of the winning assembly (# reads, #contigs, # orfs). MaSuRCA did not run on this sample because it requires paired-end input.

**Figure 4.**
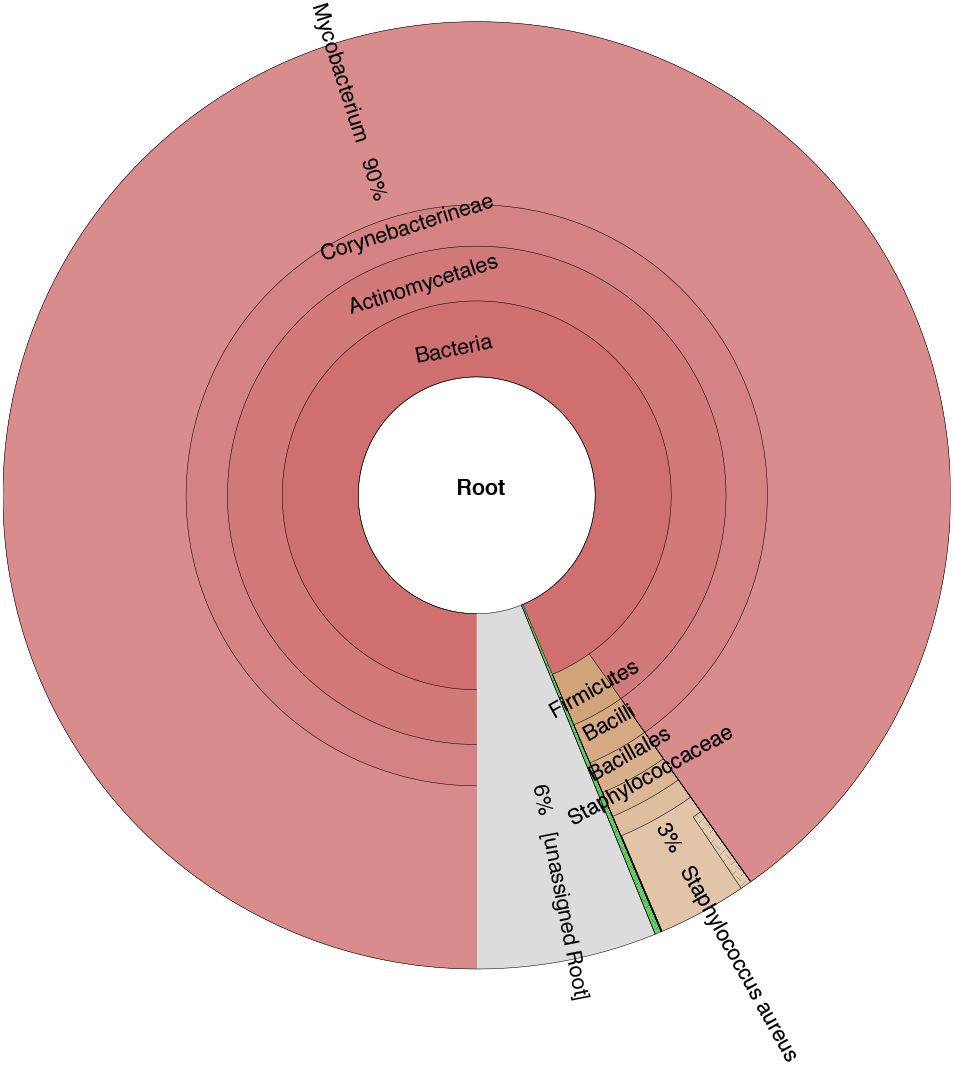
iMetAMOS classification output identifies possible contamination. On sample ERR233356 retrieved from the Sequence Read Archive, the majority of data is clearly sourced from a Mycobacterium. However, a significant fraction of the data (∼10% of reads or ∼29% of assembly) belongs to other, mostly unidentified, organisms. A subset of 3% of the reads (1.81 Mbp of the assembly) is identified as *S. aureus* and covers over 60% of the *S. aureus* genome. iMetAMOS automatically identified this potential contaminant and binned the contigs by genus to facilitate easy confirmation and removal by the user.

## Discussion

We have developed an open-source microbial analysis pipeline, iMetAMOS, which automates the process of ensemble assembly. In addition, its modular architecture is extensible and able to incorporate additional analyses or alternative tools. A potential enhancement is assembly correction, or contig breaking, which iMetAMOS does not currently support. However, the infrastructure required to support this is largely in place. For example, REAPR [12] is included with iMetAMOS and capable of splitting assembled contigs at predicted mis-assemblies. Using this and other supplied validation tools, assembly breaking could be iteratively performed until the validation scores are no longer improving or no more corrections are possible. Alternatively, because iMetAMOS generates multiple assemblies, assembly reconciliation techniques [61–63] could be incorporated into the pipeline. However, in practice, we have found the simple process of running multiple assemblers with multiple parameters is capable of generating high-confidence assemblies on its own, while merging assemblies can increase the risk of mis-assembly without significantly improving continuity [8].

The iMetAMOS extensible framework also supports customizable workflows and parameters on a per-user or per-run basis. Because all components of iMetAMOS are open source, users and tool authors are able to contribute improved parameters to the repository. Users can also contribute custom workflows tailored for specific analyses. In this way, iMetAMOS can serve as a best-practice repository for multiple assemblers, data types, and analysis tools.

## Conclusions

iMetAMOS enables accurate and reproducible genome assembly via a “GAGE-in-a-box” analysis, allowing non-expert users to run multiple assemblers, validation metrics, and annotations with a single command. Results are presented in a simplified and interactive HTML5 format, and reproducibility is enabled through detailed logging and workflows. The current implementation supports over thirteen assemblers and seven validation tools, and its modular architecture supports the easy addition of future tools. Ensemble assembly is more robust, reproducible, and accurate than manual assembly, even surpassing the quality of GAGE-B assemblies using the same data and tools. Most importantly, iMetAMOS provides users with a simple means to generate multiple assemblies and validation metrics, empowering them to choose the best assembly for their specific needs.

## Availability and Requirements

iMetAMOS is open source under the Perl Artistic License [64]. Source code and binaries are freely available from https://github.com/marbl/metAMOS. iMetAMOS is implemented in Python, C++, Perl, and Java and supported on Linux and OSX. All assemblies described here are available for download from http://www.cbcb.umd.edu/software/imetamos.

## Authors’ contributions

SK and AMP conceived the method and drafted the manuscript. SK, TJT, and CMH developed the software. MP led the development of the MetAMOS framework. All authors edited and approved the final manuscript.

## Acknowledgements

We thank Magoc *et al.* and Comas *et al.* who submitted the raw data that was used in this study. The contributions of SK, TJT, and AMP were funded under Agreement No. HSHQDC-07-C-00020 awarded by the Department of Homeland Security Science and Technology Directorate (DHS/S&T) for the management and operation of the National Biodefense Analysis and Countermeasures Center (NBACC), a Federally Funded Research and Development Center. The views and conclusions contained in this document are those of the authors and should not be interpreted as necessarily representing the official policies, either expressed or implied, of the U.S. Department of Homeland Security. In no event shall the DHS, NBACC, or Battelle National Biodefense Institute (BNBI) have any responsibility or liability for any use, misuse, inability to use, or reliance upon the information contained herein. The Department of Homeland Security does not endorse any products or commercial services mentioned in this publication. MP and CMH were supported by NIH grant R01-AI-100947and the NSF grant IIS-1117247.

